# Topology changes of the regenerating *Hydra* define actin nematic defects as mechanical organizers of morphogenesis

**DOI:** 10.1101/2024.04.07.588499

**Authors:** Yamini Ravichandran, Matthias Vogg, Karsten Kruse, Daniel JG Pearce, Aurélien Roux

## Abstract

*Hydra* is named after the mythological animal for its regenerative capabilities, but contrary to its mythological counterpart, it only regenerates one head when cut. Here we show that soft compression of head regenerating tissues induces the regeneration of viable, two headed animals. Topological defects in the supracellular nematic organization of actin were previously correlated with the new head regeneration site^1^. Soft compression creates new topological defects associated with additional heads. To test the necessity of topological defects in head regeneration, we changed the topology of the tissue. By compressing the head regenerating tissues along their body axis, topological defects of the foot and of the regenerating head fused together, forming a toroid with no defects. Perfectly ordered toroids did not regenerate over eight days and eventually disintegrated. Spheroids made from excised body column tissue partially lose their actin order during regeneration. Compression of spheroids generated toroids with actin defects. These tissues regenerated into toroidal animals with functional head and foot, and a bifurcated body. Our results show that topological defects in the actin order are necessary to shape the head of the regenerating *Hydra,* supporting the notion that actin topological defects are mechanical organizers of morphogenesis.

---

Morphogenesis and regeneration share common principles of development that establish body axes and cell fates. But less is known about how cells and tissues coordinate forces that shape organisms. *Hydra* is an established animal model for regeneration and developmental biology since the eighteenth century^2^. The evolutionarily conserved Wnt family proteins specify the body axis in the animal by forming the head organiser complex^3–6^. Upregulation of β-catenin - a mechanosensing transducer of the Wnt pathway - leads to the formation of additional heads during regeneration, showing that head determination and tissue mechanics are coupled^7^.

Mechanical cues such as osmotic oscillations are also essential for *Hydra* regeneration^8^. Breaking of symmetry in these oscillations driven by muscular contractions plays a role in specifying the body axis^9,10^. These contractions rely on ordered basal actin bundles that display supracellular organization, reminiscent of a liquid crystal-like nematic order: in the adult animal, they align parallel to the body axis in the ectoderm, forming aster-like topological defects at the foot and at the mouth, and in the endoderm, they form concentric circles around the body of the animal^1,11^. Topological defects are singularities in the nematic order, surrounded by a specific pattern of order function of their topological charge. On surfaces, the distortion of the nematic field around defects is energetically coupled to curvature, an effect that can passively drive surface deformation^12^. In living systems that behave as active materials able to generate internal forces, topological defects can generate unique patterns of stress around them which can drive motion or deformation^12^. Examples of defects acting as force-organizing centres include defects required for cell extrusion, cell sorting and collective cell migration^13^. More recently we showed that the three-dimensional growth of cellular vortices is shaped by integer topological defects^14^, a process similar to the germ band extension in drosophila^15^. These studies highlight the possible role of defects as mechanical organizers of morphogenesis in biological systems^15^. This concept is most supported by recent studies conducted in *Hydra* showing a strong correlation between the position of topological defects in the actin with the position of the new mouth and foot during regeneration^1^.

Given the nematic properties of actin^1^, we aimed at changing the nematic order in the regenerating *Hydra* to probe the role of actin defects. For this, we confined *Hydra* during head regeneration: adult animals were sectioned in their transverse plane, and the part containing the foot, hereafter called the head-regenerating tissue, was left to recover for 6 hours. Head-regenerating tissues were then subjected to soft compression between an agarose slab of varying stiffness (0.5%, 1% and 2% agarose) and a glass bottomed dish. Head-regenerating tissues were kept under compression for four days post-dissection (dpd), and then released on day five (Fig. 1a). Animals were screened at 12dpd to allow all tissues to complete regeneration before phenotypic analysis. As a control, to confirm that agarose did not impact regeneration, head-regenerating tissues were embedded in agarose. These tissues regenerated into viable uniaxial animals within 4dpd (Fig. 1b and Supplementary Video 1).

**Figure 1.**
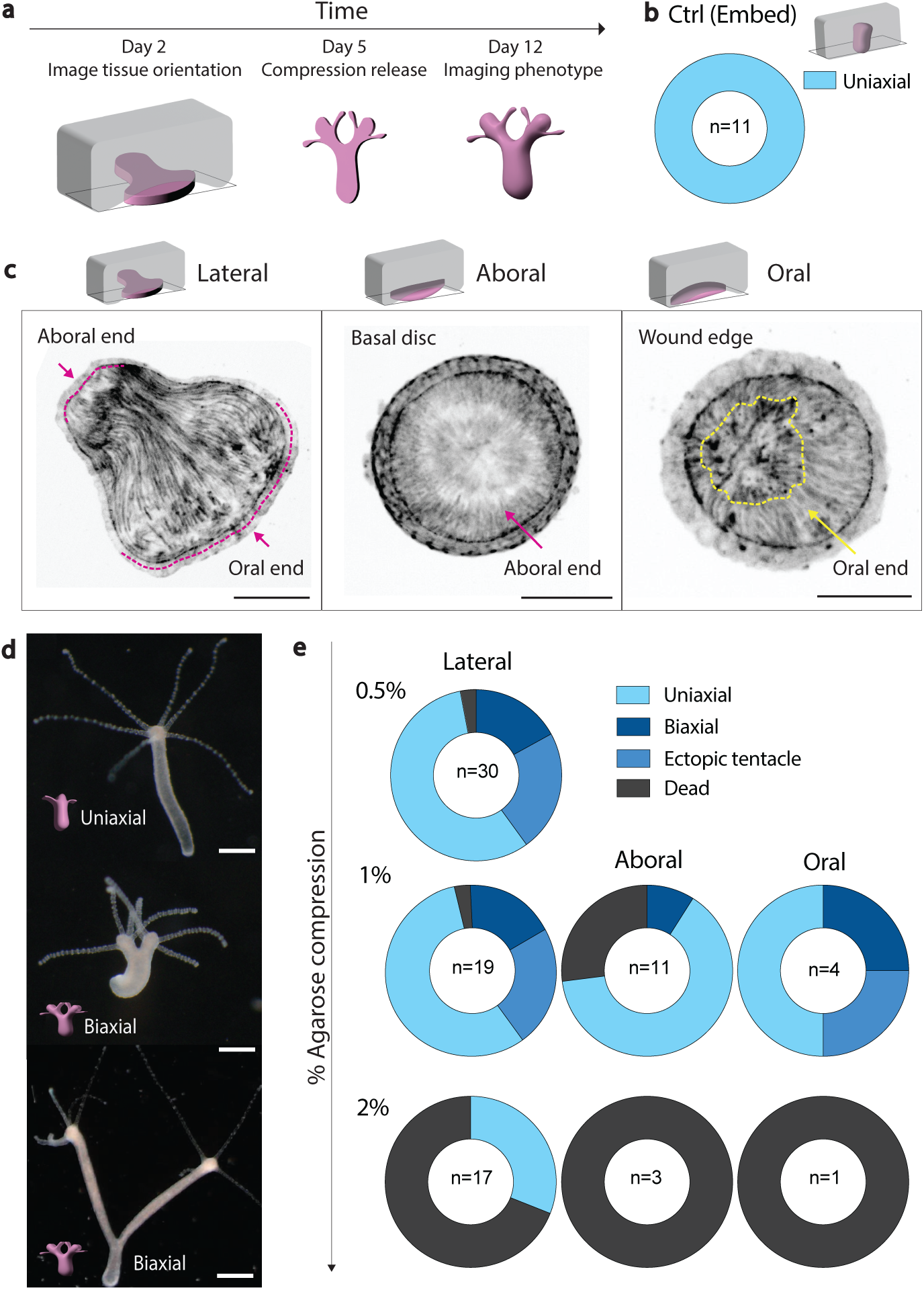
Mechanical induction of bicephalous morphogenesis in *Hydra*. **a**, Schematics showing the experimental timeline of head-regenerating tissues compression, release and phenotype analysis. **b,** Ring p showing phenotype distribution for control head regeneration experiment (0.5% agar embedded). **c,** Schematics of head-regenerating tissue orientations under compression and corresponding spinning disk microscopy images of GFP-Ecto-LifeAct tissues. Scale bars, 100 μm. Pink dashed lines, tissue contours; arrows oral/aboral ends; orange dashed line, disordered actin region. **d,** Binocular images of fully regenerated *Hydra* post compression release at 12dpd. Scale bars, 500 μm. **e,** Ring plot showing the distribution of phenotypes at 12dpd post compression release for different percentages of agar, and different orientations.

Under compression, head-regenerating tissues adopted one of three possible orientations; lateral, oral or aboral. In lateral, the tissue axis is perpendicular to the compression axis, oral corresponds to compression aligned with the tissue axis, the foot facing the glass, and aboral is the inverse of oral orientation (Fig. 1c). We captured the tissue orientation 12h post-compression (24 hours post-dissection (hpd)). We observed that under the softest 0.5% agarose compression (0.5% AC), all head-regenerating tissues oriented laterally (Extended Data Fig.1a), while with stiffer agarose (1% and 2% AC) a significant fraction displayed oral and aboral orientations indicating their inability to reorient under compression (Extended Data Fig.1a).

At 12 dpd, the most striking phenotype was a substantial proportion of bicephalous animals (30% at 0.5% AC) (Fig. 1d,e). As the single foot is symmetrically connected to the two heads, we refer to them hereafter as biaxial animals. Uniaxial animals with ectopic tentacles were also observed (25% at 0.5% AC), similar to the ectopic tentacles observed in weak Wnt3 overexpressing mutants (Extended data Fig. 1b)^6,16^. However, biaxial animals are not equivalent to Wnt3 overexpressing mutants with ectopic heads, as ectopic heads form after the establishment of the primary body axis marked with the primary head^6,16^. The biaxial animals were viable, and the two heads functional as evidenced by an *Artemia* feeding assay (Extended Data Fig. 1c and Supplementary Video 1). *In situ* hybridization of Wnt3 RNA confirmed the full differentiation of the additional heads (Extended Data Fig. 1d).

We observed that deformation of head-regenerating tissues increased gradually with the agarose stiffness (Extended Data Fig. 1e). The most notable change induced by increasing stiffness was a significant fraction of dead animals at 1% and 2% AC (Fig. 1e). Furthermore, the initial orientation dramatically changed the relative proportions of the phenotypes: tissues oriented laterally displayed the same phenotypic distribution in regenerated animals for both 0.5% and 1% AC (Fig. 1e). However, head-regenerating tissues with oral and aboral orientations, present only in 1% and 2% AC, had a much higher proportion of death (Supplementary Video 2). Thus, orientation of head-regenerating tissues impacted the phenotypic distribution more than increasing agarose stiffness (Fig. 1e).

To test whether soft compression was essential for phenotype generation, we compressed the head- regenerating tissues between stiffer plastic microfluidic channels of fixed thickness. At 200 microns confinement, only dead tissues were obtained, and at 400 microns, no phenotypic changes were observed (Extended Data Fig. 1f,g). Moreover, compressing head-regenerating tissues between two 1% agarose slabs increased the penetrance of phenotypes, while reducing the proportion of deaths (Extended Data Fig.1h). We therefore concluded that soft compression was essential, probably applying sufficient constraints to cause biaxial phenotype, still allowing for contractions essential for regeneration^8^. Our results show that soft compression of foot tissues during regeneration of the head can trigger head duplication, leading to viable bicephalous animals.

We further investigated the reasons for such a dramatic change in morphogenetic outcome. Given the correlation between the position of actin topological defects and site of new head regeneration^1^, we wondered how the actin nematic order was modified during compression. 3D two photon imaging of biaxial animals (12dpd) showed that ectodermal actin retains its long-range nematic order, but that additional defects were present at the mouth position, and in between the heads (Extended Data Fig. 2a). We wondered when these additional defects appeared under compression.

To visualize actin order during regeneration, head-regenerating tissues were prepared from Lifeact-GFP expressing animals^17^ and imaged under compression by live spinning disk confocal microscopy. Head-regenerating tissues inherited the actin nematic order, with a single topological defect on the basal disc, and longitudinal fibres expanding towards the regenerating wound (Fig 1a, 2a). Upon compression the wound edge was strikingly flattened out and regenerated two heads at the tissue extremities in 30% of the 0.5% AC animals within 4dpd (Fig 2b, Supplementary Video 2). In animals that successfully regenerated, actin order was retained throughout regeneration confirming that long range actin order was mostly kept during compression (Fig 2a).

**Figure 2.**
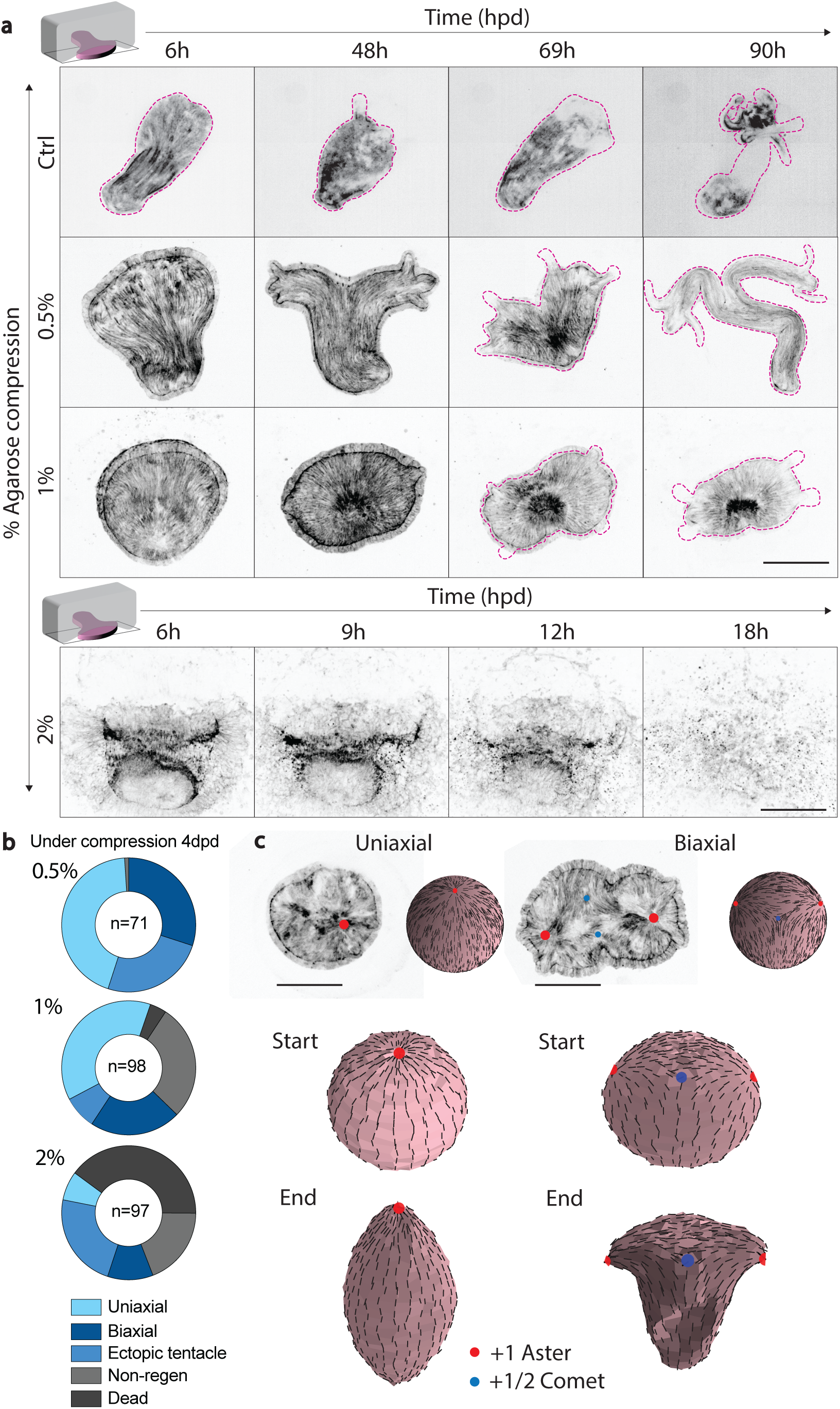
Regenerating tissues form two heads and two aster defects during compression. **a**, Montages of live spinning disk microscopy videos (maximum intensity projections) of GFP-Ecto-LifeAct head-regenerating tissues for different % agar compression. Control is 0.5% agar embedded (Supplementary Video 2-5). Pink dashed lines correspond to tissue contours. Scale bars, 100 μm. **b,** Ring plot showing phenotype distribution at 4dpd under different % agar compression. **c,** Simulations of deformation of active nematic spheres. Upper row, initial state of the nematic order with corresponding picture of a head-regenerating tissue. Left column, uniaxial, with 2 aster defects (red dot, the second - symmetrical - is not visible); right column, biaxial with 3 aster defects (red dots, one is not visible) and compensating -1/2 defects (blue dots). Middle row, shape of the active nematic sphere at the beginning of simulations. Bottom raw, shapes of active nematic spheres at the end of simulations. Scale bars 100 μm.

During regeneration under 0.5% AC, proportions of each phenotype were similar than observed previously in the fully regenerated animals (Fig 2b). In less than 1% of the cases, tissues did not regenerate but remained viable until the end of compression (4dpd) (Fig 2a,b). The non-regenerative fraction significantly increased in 1% AC (28%), as higher stiffness may slow down the tissues’ contractions (Supplementary Video 3) delaying regeneration, and death became significant. For 2% AC, 40% of the head-regenerating tissues underwent death, while uniaxial tissues became minor. Thus, the phenotypes appeared during compression, and increasing agarose stiffness increased prevalence of mechanically induced phenotype and death. Therefore, phenotypes visible in the released animals appeared earlier because of compression (Fig 1e, Supplementary Video 4).

In 1% and 2% AC, some of the head-regenerating tissues regenerating two heads were orally oriented, with the mouth/wound side facing the glass. In this case, two +1 aster defects were clearly visualized while the two heads emerged (Fig. 2c, Extended Data Fig.2b), consistently with the reported correlation between +1 aster defect position and head regeneration^1^. We concluded from these results that the regeneration of two heads correlated with the emergence of two aster defects under compression. Based on our previous findings that integer topological defects can organize cellular stresses that shape tissues we imagined that the tissue shape required for regenerating the head could be generated by the actin stress field around the defects^14^. We further envisioned that a second defect could create a tissue shape adapted to a second head.

To test these hypotheses, we undertook a theoretical approach. We approximate the head-regenerating tissue as a thin elastic nematic material in which active stress is generated parallel to the nematic orientation. We approximate the initial shape of the regenerating *Hydra* as a sphere and the actin supracellular organization is described by the nematic orientation field on its surface.

We assume that all nematic orientation fields feature an aster at the pole corresponding to the foot. The remaining orientation field is broadly inferred from experimental images. For uniaxial regeneration, the actin supracellular organisation features a single aster on the mouth pole (Fig 2c). At the end of uniaxial simulations, the active elastic adopts an elongated shape similar to a single headed *Hydra*, with defects at the tips (Fig 2c, Supplementary Video 5). In a thin elastic active nematic, aster defects are able to generate protrusions by organizing active stress^18,19^. During biaxial regeneration of *Hydra*, we observed a third aster associated with the additional head (Fig. 2c), which we included in the initial nematic orientation field; this required an additional pair of negative defects to preserve the total topological charge. During biaxial simulations, the spherical surface undergoes a splitting of the central axis, resulting in a bi-axial shape (Fig 2c, Supplementary Video 6). Thus, our simulations show that the shape of a second head can emerge from an additional aster in the orientation field of the supracellular actin. Our results show that each head regeneration correlates with an aster defect of the actin supracellular organization on the head-regenerating tissue, supporting the notion that an actin defect is required to shape a head.

To test further the requirement of actin-defects, we next turned our attention to head-regenerating tissues that failed to regenerate under compression, previously referred to as non-regenerative tissues. These may be just delayed in regeneration or may lack features essential to regeneration. To test whether compression had simply slowed down regeneration, we determined the ability of 1% AC non-regenerative tissues to regenerate post compression release. These tissues were released from compression at 5dpd and a large proportion of these non-regenerative tissues did regenerate as screened at 12dpd (> 80%, Fig. 3a). The remaining 20% (approx. 5% of the initial pool) were unable to regenerate even after release. We defined them as persistent non-regenerative tissues (Fig. 3b). Live colour imaging 12h post release showed the presence of a longitudinal tube-like thickening at the centre of the persistent non-regenerative tissues (Fig. 3c, Extended Data 3a). Epifluorescence imaging revealed the presence of a tissue fold forming a tunnel through the tissue resulting in the topology of a torus (Fig. 3b). By comparison at 12h post release, tissues with a spherical topology had already regenerated primitive tentacles (Fig. 3c). 360° light-sheet microscopy confirmed the toroidal topology of the persistent non-regenerative tissues, hereafter called toroids. These images also showed that the actin orientation field displayed rotational symmetry, thus featuring no defects (Fig. 3d and Extended Data Fig. 3b). As its Euler characteristic equals zero, the torus is the only topology supporting a fully ordered actin superstructure, with no defects. The lack of defects in toroids could therefore be associated with their inability to regenerate.

**Figure 3.**
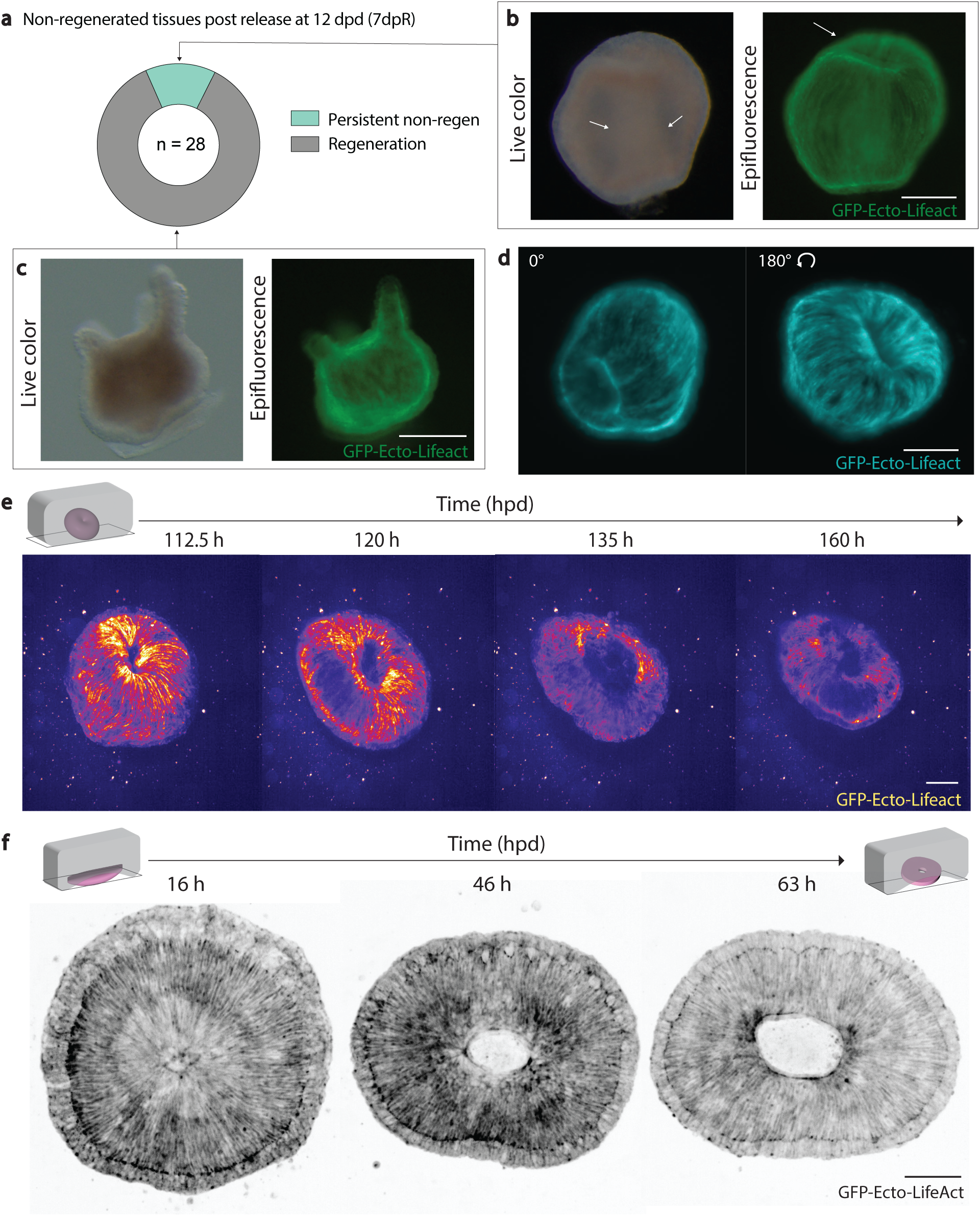
Defect-less toroids fail to regenerate. **a**, Ring plot showing fate of non-regenerated tissues after 4dpd compression, and screened at 12dpd (5 days post-release). **b,** Left, live colour image of persistent non-regenerative tissues 12h post compression release; white arrows show tubular tissue thickening. Right, corresponding epifluorescence image of the tissue expressing GFP-Ecto-Lifeact displaying a tissue fold at the top (white arrow). Scale bars, 500 μm. **c,** Left, live colour image of regenerated tissues 12h post compression release (non-regenerated after 4dpd compression) displaying tentacles. Right, corresponding epifluorescence image of the tissue expressing GFP-Ecto-Lifeact. Scale bars, 200 μm. **d,** Maximum intensity projections of a light-sheet microscopy stack of GFP- Ecto-Lifeact expressing *Hydra* toroid. Left, top view. Right, bottom view. Scale bar 100 μm. **e,** Spinning disk microscopy timelapse (maximum intensity projections) of GFP-Ecto-LifeAct expressing *Hydra* toroid embedded in 0.5% agarose. Fire LUT was applied to appreciate three dimensionality. Scale bar, 100 μm. **f,** Spinning disk microscopy timelapse (maximum intensity projections) of GFP-Ecto-Lifeact head-regenerating tissue forming a toroid. Scale bar, 100 μm.

To confirm the lack of regeneration after release, toroids were imaged live. The toroids retained their shape over 60 hrs post release and did not break symmetry in their actin order nor regenerate (Fig 3e, Supplementary Video 7). The toroids eventually disintegrated before 12dpd, explaining why only dead or regenerated animals were observed at 12dpd (Fig 1c). In conclusion, head- regenerating tissues that acquired the topology of a torus, with no topological defects in the actin superstructure, could not regenerate.

We then studied how this unique topology change could occur under compression. In aboral and oral orientations, head-regenerating tissues occasionally underwent tissue tear right at the centre of the actin aster suggesting a local increase in stress (Fig 3f). A mirror tear occurred at the antipole of the head-regenerating tissue. The tissue tears during contractions, which are oriented radially around the defect (Supplementary Video 8). Wound healing occurs along the body axis fusing the two tears, annihilating the two aster defects, and giving rise to a defect-less toroid (Fig 3f)^20^.

As mentioned previously, aster defects deform thin active nematic elastic materials into a protrusion, and the compression against the defect could invert the protrusion^18^. As a consequence, the two ends of non-regenerative tissue would buckle inward under axial compression, which only occurs for oral or aboral orientations. To test this, we initialize our simulations with two asters placed at opposite poles and compression along the same axis. Simulations show that compression indeed inverts the protrusions that result from aster defects (Fig. 4a). In our experiments, we observed the same inversion of protrusions associated with asters at foot sites under compression (Fig. 4b Extended Data Fig. 4c). This simultaneous inward buckling of tissue defects aids in the fusion event required for toroid formation (Fig. 3f), while preserving the symmetry allowing for a defect-less actin superstructure.

**Figure 4.**
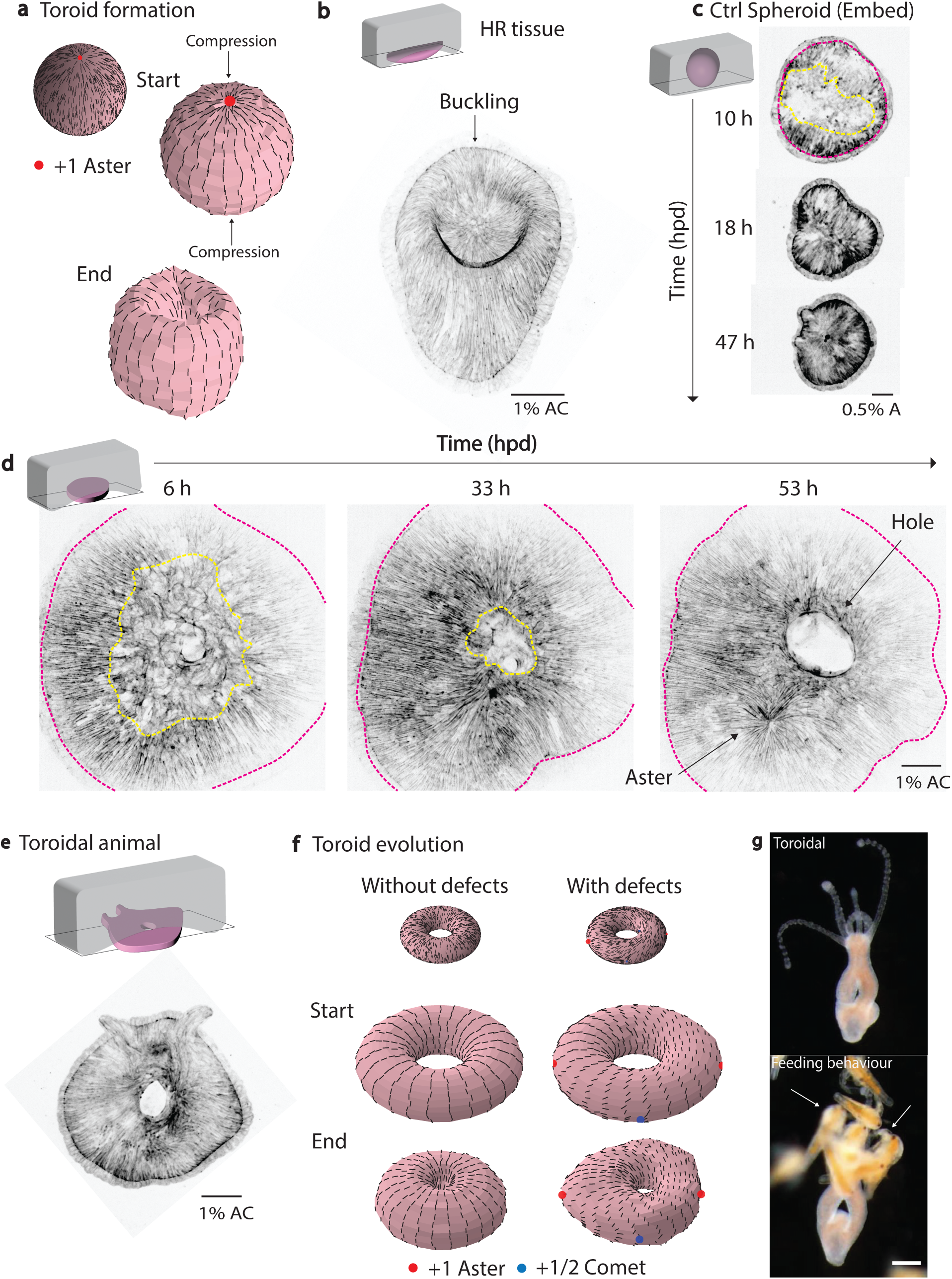
Actin topological defects are necessary to shape head regeneration in *Hydra* toroids. **a**, Simulation of toroid formation under compression. Top, initial order of the active nematic sphere, with corresponding Head-regenerating tissue image. Middle, shape of the sphere at the start of simulations, arrows indicate orientation of the constrain. Bottom, shape of the active nematic sphere at the end of the simulation. **b,** Head-regenerating tissue under compression displaying buckling at +1 defect site. **c,** Control experiment of spheroid tissue regenerating in embedded 0.5% agarose. Pink dash line shows tissue contours, and orange dash line shows the area where actin lost nematic order. **d,** Montage of spheroid tissue under 1% agar compression undergoing tissue tear and forming +1 aster defect (arrow). Pink dash line shows tissue contours, and orange dash line shows the area where actin lost nematic order. **e,** Toroidal tissue under compression regenerating a head with tentacles at 84hpd. **f,** Simulations of toroid evolution, with and without defects. **a-e,** Scale bars, 100 μm. **g,** Live colour images of **top,** toroidal animal and **bottom,** feeding behaviour of toroidal animal. Scale bar, 200 μm

Changes in topology are non-continuous changes in a shape: they require tearing and/or fusion of surfaces and are only witnessed during drastic stages of morphogenesis such as gastrulation. To discriminate whether the change of topology or the lack of defects blocks regeneration, we aimed to generate toroids with topological defects and test whether they would regenerate. Previous work^1,21^ using the excised body column tissue of *Hydra* reported partial loss of supracellular actin order during initial stages of regeneration while it reshapes as a sphere, referred to as spheroid hereafter (Fig. 4c). Spheroids have been shown to regenerate into both uniaxial and biaxial animals (Extended Data Fig. 4a)^21^, suggesting that spheroids can have additional defects, generating new heads. We hypothesized that when subjected to compression, a fraction of spheroids will undergo tear and form toroidal tissues featuring additional topological defects.

Spheroids under compression were imaged by live spinning-disk confocal microscopy. Large areas where actin lacked nematic order were visible (Fig. 4c). Initially, no topological defects were seen, but 16hpd, the actin superstructure emerged and topological defects appeared (Fig. 4c Time 10- 36hpd, Supplementary Video 10). As expected, certain tissues underwent tear and changed topology before global actin order was established (Fig. 4d). This is opposed to the topological change of toroids reported above, which occurs with preserved actin order. Furthermore, aster defects were clearly visible in the actin superstructure of spheroids when the tissue was still under compression undergoing wound healing (Fig. 4d, Time 53hpd, Supplementary Video 11). At the position of these defects, head and foot features appeared, showing that toroidal tissues with defects are able to regenerate (Fig. 4e).

Among the regeneration features that appeared on toroidal tissues with defects were protuberances on their contour, reminiscent of dome shapes coupled to aster defects (Fig. 4d,e)^18^. To test whether these protuberances could be created by stress gradients linked to defects, we simulated toroidal surfaces with and without defects (Fig. 4f). Simulations of toroids with no defects fail to break symmetry and remains perfectly toroidal (Supplementary Video 12), whereas simulations featuring defects break symmetry and form protuberances colocalizing with defects (Supplementary Video 13).

To test whether toroids with defects would further regenerate into viable animals, we released them from compression at 5dpd and observed that they were able to recover and thrive. After compression, we observed a multitude of phenotypes including toroidal animals with a head and foot (Fig. 4f), and new phenotypes such as bipedal animals (Extended Data Fig. 4a). The induction of tissue tear and consequent wound healing depended on the stiffness of agarose used for compression, as toroidal animals were only generated using 1% AC (Extended Data Fig. 4c). These animals have fully functional head and foot, able to trap *Artemia* (Supplementary Video 14), while having a hole that bifurcates their gastrovascular cavity (Fig. 4g). Overall, *Hydra* toroidal tissues with actin defects regenerate live toroidal animals with a head and a foot, conserving their unique topology, showing that actin topological defects are necessary for forming a new head and establishing body axis.

Overall, these results strongly support a necessary role of actin integer topological defects in the regeneration of *Hydra’s* head. Simulations based on active nematics theory support that these actin defects establish the stress field required for positively curved surfaces shaping the new head. These simulations are supported by the observations that tearing and buckling occur predominantly at the defects, as expected from the stress field calculated from the theory. How the mechanical role of actin topological defects is coupled to the Wnt organizer remains to be explored. However, recent findings propose that stretching of cells associated with larger stresses at the defects may be coupled to Wnt production through mechano-sensing signalling^22^ in line with past findings suggesting a tight coupling mechanochemical coupling in the animal^8,9,23^.

The common topology of developing and regenerating tissues is equivalent to the one of a sphere, with a total charge of +2. Our work shows that topology is an important constraint that participate in the establishment of the body axis and thus the body plan. Topology thus may have participated in the selection of common features of body plans through evolution. It is also notable that embryos change their topology at critical steps during their development: during gastrulation, but also during somitogenesis or neurulation. Thus, genetically controlled changes in the embryo topology may participate in establishing the final body plan. Our work opens new perspectives for understanding how topological changes are required for morphogenesis, but also on how topology could be used to create new body plans.

## Supporting information

Supplementary Figures & Theory

## Movie captions

**Movie 1:** Biaxial animal feeding behaviour

**Movie 2:** Ctrl 0.5% agarose embedded animal regenerating one head

**Movie 3:** 0.5% agarose compressed head-regenerating tissue displaying biaxial phenotype post regeneration

**Movie 4:** 1% agarose compressed head-regenerating tissue displaying biaxial phenotype post regeneration

**Movie 5:** 2% agarose compressed head-regenerating tissue dying by disintegration

**Movie 6:** Simulation of uniaxial spheroid under compression

**Movie 7:** Simulation of biaxial spheroid under compression

**Movie 8:** Spinning disk video of toroid non regenerating post release

**Movie 9:** Toroid formation under agarose compression

**Movie 10:** Simulation of uniaxial spheroid under compression

**Movie 11:** Regeneration of an embedded spheroid (control)

**Movie 12:** Regeneration of a toroidal tissue with aster defects into toroidal animal.

**Movie 13:** Simulation of defectless toroid retains rotational symmetry

**Movie 14:** Simulation of toroid with defects generating protuberances correlated with defect regions

**Movie 15:** Toroidal animal feeding behaviour

All videos have been compressed to 10fps unless specified in movie filename and playback speed is 1x

## Methods

### Animal culture

All of the experiments were performed using transgenic *Hydra vulgaris* (strain Basel) expressing Lifeact-GFP in the ectodermal cells provided by B. Hobmayer from the University of Innsbruck, Austria^11^.

Cultures were maintained in *Hydra* Medium (HM: 1 mM NaCl, 1 mM CaCl2, 0.1 mM KCl, 0.1 mM MgSO4, 1 mM Tris pH 7.6) at 18 °C. Animals were fed two to three times per week with freshly hatched *Artemia nauplii* and starved for 24h before any experiment. Non-budding animals that had fed were chosen at random from the dish.

### Sample preparation

Animals approximately greater than 5mm long were transversely sectioned into three equal parts. The head tissue was discarded. The body column was left to heal for 6h to form sealed spheroids. The bottom third containing the foot was left to heal for 6h and used as the head-regenerating tissue. A scalpel equipped with a no. 23 blade was used for dissections.

For control experiments, spheroids and head-regenerating tissues were embedded in a soft gel (0.5% low-melting-point agarose (Sigma) prepared in HM)^1^. The regenerating tissues were placed in cooled down liquefied gel 6h post dissection and the gel were left to solidify. The tissues naturally settled down to the glass bottom due to their density which made it easier to image them.

### Compression experiments

20 ml of liquefied agarose (0.5%, 1% or 2%) was poured in a 10 ml petri dish in order to obtain a constant height for the agarose slabs. The agarose was allowed to solidify with closed lids to prevent evaporation. Once cooled, regenerating tissues were placed on the solidified agarose. With the tissue at the centre an approximate 5mm x 5mm square cut is made in the agarose with a no.23 scalpel. Then, the agarose slab containing the tissue on its surface is gently excised out of the petri dish with the scalpel and flipped over tissue side first on a 35mm glass bottomed Matek dish, trapping the regenerating tissues between the glass bottom and the excised agarose slab. Extreme precision is required during this step to avoid tissue shear and subsequent disintegration. After placing roughly 4-5 agarose compressed tissues in each Matek, 3ml of liquefied agarose is poured on top, in order to avoid evaporation and preventing tissues wiggling their way out of the compression. After the agarose has been solidified, 2 ml of HM is added to each petri dish and lids placed to further prevent evaporation during the course of the 4 days of regeneration at 18°C.

For agarose/agarose compression experiments, the Matek is coated with 1ml of 1% agarose prior to compressing tissues with 1% agarose slabs.

### Compression release

On 5dpd the supernatant HM is removed from each dish. 5mm x 5mm cuts are made in agarose around the tissues in the centre, located at the bottom. HM is flushed into the cuts which enables the agarose slab to float and with it the compressed tissue at the bottom. This enables us to minimize any shear faced by the tissue during compression release. The suspended tissue/regenerated animal is collected with a glass Pasteur pipette and moved to a new well with fresh HM.

For experiments where phenotypic screening was performed all the tissues were labelled on the Matek with corresponding numbers and when released placed in labelled wells. This animal labelling enabled us to identify the correlation between tissue orientation and what they regenerated into when their phenotype was screened upon compression release.

### Microfluidic channel confinement

200 um and 400 um ibidi sticky slide were used for channel confinement experiments. The tissue was placed in the sticky channel and excess water removed then sealed with a glass coverslip. Hydra media was then flushed into the wells gently and replenished every day to compensate for evaporation loss. The tissues were confined for 4 days before images were taken to screen for phenotype.

### Microscopy

All time-lapse imaging was performed with an inverted microscope Nikon Ti-E installed in a room where temperature was maintained at 20°C. The microscope was also equipped with an automated stage and a Yokogawa CSU-W1 spinning disk unit. Image acquisition was performed with an Andor Zyla 4.2 Plus camera, operated with Slidebook Software. Fluorescence 4D time-lapse imaging was performed to capture actin dynamics in regenerating tissues either embedded or under agarose slab compression using a10x (NA 0.30) objective. For all experiments under compression, we acquired 2 images/h for 84h. All videos are maximum intensity z-projections of 42 z-stacks each spaced 4um each.

Light-sheet microscopy was performed on a Miltenyi Biotec Ultramicroscope Blaze Light Sheet equipped with 4.2 Megapixel sCMOS camera. 4X (NA 0.35) objective was used with 2.5x zoom. The toroid was embedded in an agarose cube with HM buffer, and imaged in the light sheet. The agarose cube was manually rotated to obtain different angles of the tissue. 4um thickness stacks were taken. 184 z-stacks were taken and maximum intensity z-projected to compile final images.

An upright multiphoton confocal microscope - Leica SP8DIVE FALCON equipped with HyD detectors was used to obtain the 3D images of bicephalous animals. Live imaging was possible and ability to penetrate the tissue was achieved by using z-compensation mode where the laser intensity was increased with increasing depth into the tissue. Tuneable multi-photon laser was generated at 820nm to excite the GFP tagged Lifeact at 488nm. 25X (NA 0.95) IRAPO (max transmission of IR and Vis and minimal axial shift up to 1300nm) water immersion objective was used to image the samples. 4um z-stacks were used to image the samples in 3D.

### Whole mount *In Situ* Hybridization

*Hydra* at 12 dpd (7dpR) were relaxed in 2% urethane/HM for one minute, fixed in 4% PFA prepared in HM (pH 7.5) for 4h at RT and stored in MeOH at −20 °C for at least one day. Samples were rehydrated through a series of ethanol, PBSTw (Phosphate Buffer Saline, Tween 0.1%) washes (75%, 50%, 25%) for 5 min each, washed 3× with PBSTw for 5 min, digested with 10 μg/mL Proteinase K (PK, Roche) in 0.1% SDS, PBSTw for 10 min, stopped by adding Glycine (4 mg/mL) and incubated for 10 min. Samples were washed 2x in PBSTw for 5 min, treated with 0.1M TEA for 2 × 5 min, incubated 5 min after adding acetic anhydride 0.25% (v/v), 5 min after adding again acetic anhydride 0.25% (final concentration 0.5% v/v). Samples were then washed in PBSTw 2×5 min, post-fixed in 4% formaldehyde, PBSTw for 20 min, washed in PBSTw 4×5 min before adding the pre-warmed pre-hybridization buffer (PreHyb: 50% Formamide, 0.1 % CHAPS 1× Denhardt’s, 0.1 mg/mL Heparin, 0.1% Tween, 5x SSC) and incubated for 2h at 58 °C. Next, 350μL hybridization buffer (PreHyb containing 0.2 mg/mL t-RNA, 5% Dextran) containing 200 ng DIG-labelled riboprobe was heated 5 min at 80 °C, then placed on ice for 2 min. This mix was added onto the samples, then incubated for 19h at 58 °C.

Next, the samples were rinsed 3x in pre-warmed PostHyb-1 (50% formamide, 5x saline-sodium citrate (SSC)) and successively incubated for 10 min at 58°C in PostHyb-1, PostHyb-2 (75% PostHyb-1, 25% 2x SSC, 0.1% Tween), PostHyb-3 (50% PostHyb- 1, 50% 2× SSC 0.1% Tween) and PostHyb-4 (25% PostHyb-1, 75% 2× SSC, 0.1% Tween). Samples were then washed 2× 30 min in 2× SSC, 0.1% Tween, 2×30 min in 0.2x SSC, 0.1% Tween, 2× 10 min in MAB-Buffer1 (1× MAB, 0.1% Tween), blocked in MAB-Buffer2 (20% sheep serum, MAB-Buffer1) for 1h and incubated with anti-DIG-AP antibody (1:4000, Roche) in MAB-Buffer2 overnight at 4 °C.

Next, the samples were washed in MAB-Buffer1 for 4×15 min, then in NTMT buffer (NaCl 0.1 M, Tris-HCl pH 9.5 0.1 M, Tween 0.1%) for 5 min and finally in NTMT, levamisole 1 mM for 2×5 min. The colorimetric reaction was started by adding staining solution (Tris-HCl pH9.5 0.1 mM, NaCl 0.1 mM, Polyvinyl alcohol 7.8%, levamisole 1 mM) containing NBT/BCIP (Roche). The background colour was removed by a series of washes in EtOH/PBSTw (30%/70%, 50%/50%, 70%/30%, 100% EtOH, 70%/30%, 50%/50%, 30%/70%), PBSTw 2 × 10 min. Samples were post-fixed for 20 min in formaldehyde 3.7% diluted in PBSTw, washed in PBSTw 3 × 10 min and mounted with Mowiol. All steps were performed at RT unless indicated otherwise.

### Statistical analyses

All graphs were made with the GraphPad Prism7 software. Two photon images were processed using Imaris 8 software and all other images were processed using Fiji software. Schematic illustrations and figures were assembled using Adobe Illustrator 2023.

## References

1. Maroudas-Sacks, Y. et al. Topological defects in the nematic order of actin fibres as organization centres of Hydra morphogenesis. Nature Physics 2020 17:2 17, 251–259 (2020).

2. Galliot, B. Hydra, a fruitful model system for 270 years. International Journal of Developmental Biology vol. 56 411–423 Preprint at 10.1387/ijdb.120086bg (2012).

3. Hobmayer, B. et al. WNT signalling molecules act in axis formation in the diploblastic metazoan Hydra. Nature 407, 186–189 (2000).

4. Lengfeld, T. et al. Multiple Wnts are involved in Hydra organizer formation and regeneration. Dev Biol 330, 186–199 (2009).

5. Broun, M., Gee, L., Reinhardt, B. & Bode, H. R. Formation of the head organizer in hydra involves the canonical Wnt pathway. Development 132, 2907–2916 (2005).

6. Vogg, M. C. et al. An evolutionarily-conserved Wnt3/β-catenin/Sp5 feedback loop restricts head organizer activity in Hydra. doi:10.1038/s41467-018-08242-2.

7. Gee, L. et al. β-catenin plays a central role in secng up the head organizer in hydra. Dev Biol 340, 116–124 (2010).

8. Ferenc, J. et al. Mechanical oscillations orchestrate axial patterning through Wnt activation in Hydra. Sci Adv 7, 6897 (2021).

9. Kücken, M., Soriano, J., Pullarkat, P. A., Ott, A. & Nicola, E. M. An Osmoregulatory Basis for Shape Oscillations in Regenerating Hydra. Biophys J 95, 978–985 (2008).

10. Szymanski, J. R. & Yuste, R. Mapping the Whole-Body Muscle Activity of Hydra vulgaris. Current Biology 29, 1807–1817.e3 (2019).

11. Aufschnaiter, R., Wedlich-Söldner, R., Zhang, X. & Hobmayer, B. Apical and basal epitheliomuscular F-actin dynamics during Hydra bud evagination. Biol Open 6, 1137– 1148 (2017).

12. Pearce, D. J. G., Thibault, C., Chaboche, Q. & Blanch-Mercader, C. Passive defect driven morphogenesis in nematic membranes. (2023).

13. Doostmohammadi, A. & Ladoux, B. Physics of liquid crystals in cell biology. Trends Cell Biol 1–11 (2021) doi:10.1016/j.tcb.2021.09.012.

14. Guillamat, P., Blanch-Mercader, C., Pernollet, G., Kruse, K. & Roux, A. Integer topological defects organize stresses driving tissue morphogenesis. Nature Materials 2022 1–10 (2022) doi:10.1038/s41563-022-01194-5.

15. Nandi, S., Balse, A., Inamdar, M. M., Kumar, K. V. & Narasimha, M. Actomyosin cables position cell cohorts during Drosophila germband retraction by entraining their morphodynamic and mechanical properties. bioRxiv 2022.09.23.509113 (2022) doi:10.1101/2022.09.23.509113.

16. Mercker, M. et al. β -Catenin and canonical Wnts control two separate pattern formation systems in Hydra : Insights from mathematical modelling Summary of experiments contrasting the function of β -Catenin and HyWnt3. bioRxiv 1–19 (2021) doi:10.1101/2021.02.05.429954.

17. Aufschnaiter, R., Wedlich-Söldner, R., Zhang, X. & Hobmayer, B. Apical and basal epitheliomuscular F-actin dynamics during Hydra bud evagination. Biol Open 6, 1137– 1148 (2017).

18. Pearce, D. J. G., Gat, S., Livne, G., Bernheim-Groswasser, A. & Kruse, K. Defect-Driven Shape Transitions in Elastic Ac7ve Nematic Shells.

19. Duffy, D., et al. Shape programming lines of concentrated Gaussian curvature. J Appl Phys 129, 224701 (2021).

20. Lubensky, T. C. & Prost, J. Orientational order and vesicle shape. Journal de Physique II 2, 371–382 (1992).

21. Livshits, A., Shani-Zerbib, L., Maroudas-Sacks, Y., Braun, E. & Keren, K. Structural Inheritance of the Actin Cytoskeletal Organization Determines the Body Axis in Regenerating Hydra. Cell Rep 18, 1410–1421 (2017).

22. Suzuki, R. et al. Spatio-temporal Coordination of Active Deformation Forces and Wnt / Hippo-Yap Signaling in Hydra Regeneration. bioRxiv 2023.09.18.558226 (2023) doi:10.1101/2023.09.18.558226.

23. Perros, T. et al. Mechanical characterization of regenerating Hydra tissue spheres Mechanics of regenerating Hydra. bioRxiv 2023.10.16.562504 (2023) doi:10.1101/2023.10.16.562504.

