## Supplementary Figures & Theory for "Topology changes of the regenerating *Hydra* define actin nematic defects as mechanical organizers of morphogenesis"

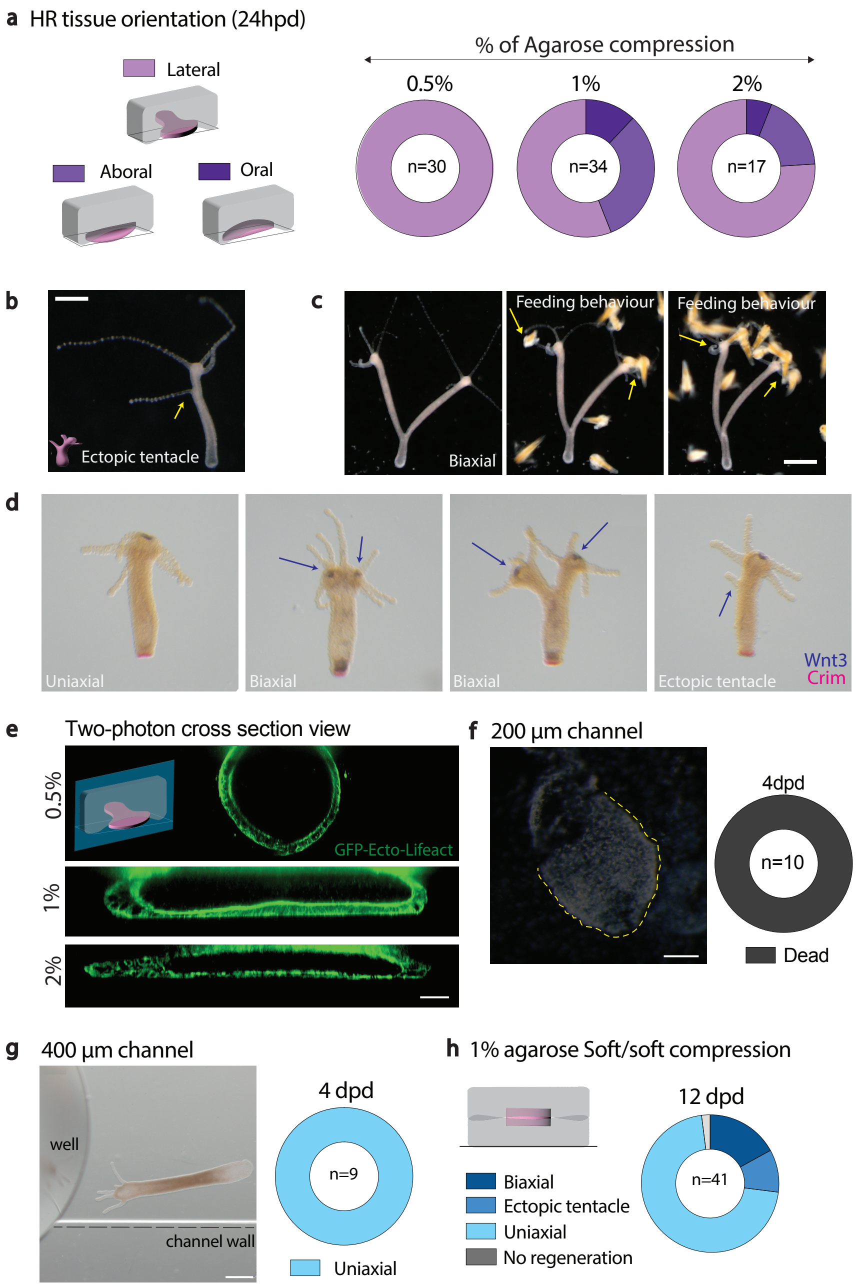


Extended Data Fig.1

**Extended Data Fig.1 a,** Donut graphs showing preferred tissue orientation at 24hpd under different percentages of AC. **b,** Live colour images of fully regenerated *Hydra* post compression release at 12dpd. **c,** Live colour images of right biaxial animal and middle and left, feeding behaviour of biaxial animal. **b-c,** Scale bars, 500 µm. **d,** *In situ* hybridization experiments for Wnt3 (head marker) and Crim (foot marker) **e,** Two-photon microscopy of the HR tissue compressed under different percentages of agarose, top panel contains schematics with a green plane showing the cross sectional view of the tissue being imaged. **f,** Live color image of tissue under 200 µm channel confinement at 4dpd and corresponding donut graph showing the phenotypic distribution. Scale bars, 200 µm. **g,** Live color image of tissue under 400 µm channel confinement at 4dpd and corresponding donut graph showing the phenotypic distribution. Scale bars, 500 µm. **h,** Schematic describing head-regenerating tissue subjected to 1% agarose soft/soft of compression and corresponding donut graph showing phenotypic distribution.

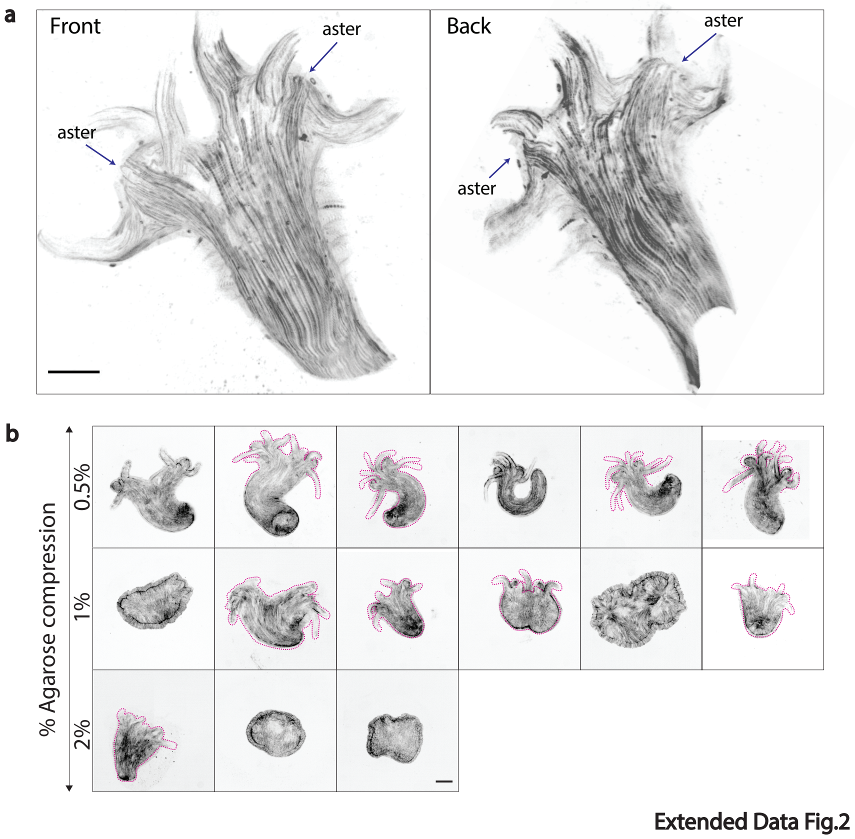


**Extended Data Fig.2 a,** Two-photon microscopy (max intensity z-proj) of biaxial *Hydra* with right, Front and left, Back view of the animal with aster topological defects marked with black arrows. Scale bar, 70µm. **b,** Live spinning disk images **(**Max intensity z-proj) at 4dpd of the various HR tissues under different % of AC displaying diversity in biaxial phenotype. Pink dashed lines correspond to tissue contours. Scale bar, 100µm.

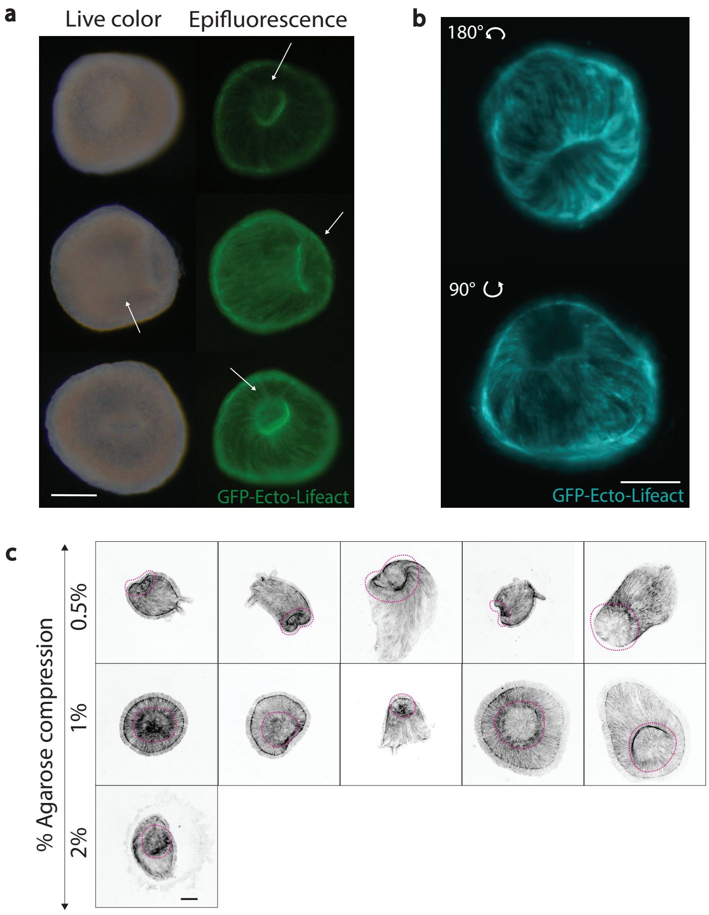


**Extended Data Fig.3 a,** Left, live colour images and corresponding right, epifluorescence images of persistent non-regen tissues with visible internal tissue thickening and tissue folds marked with white arrows. Scale bars, 200µm. **b,** Light-sheet microscopy images (max intensity z-proj) of GFP-Ecto-Lifeact expressing *Hydra* toroid. Left, angled top view. Right, angled bottom view. Scale bar 100 µm. **c,** Live spinning disk images **(**Max intensity z-proj) at 4dpd of the various HR tissues under compression with the view of buckling at the aboral end. Scale bar, 100µm.

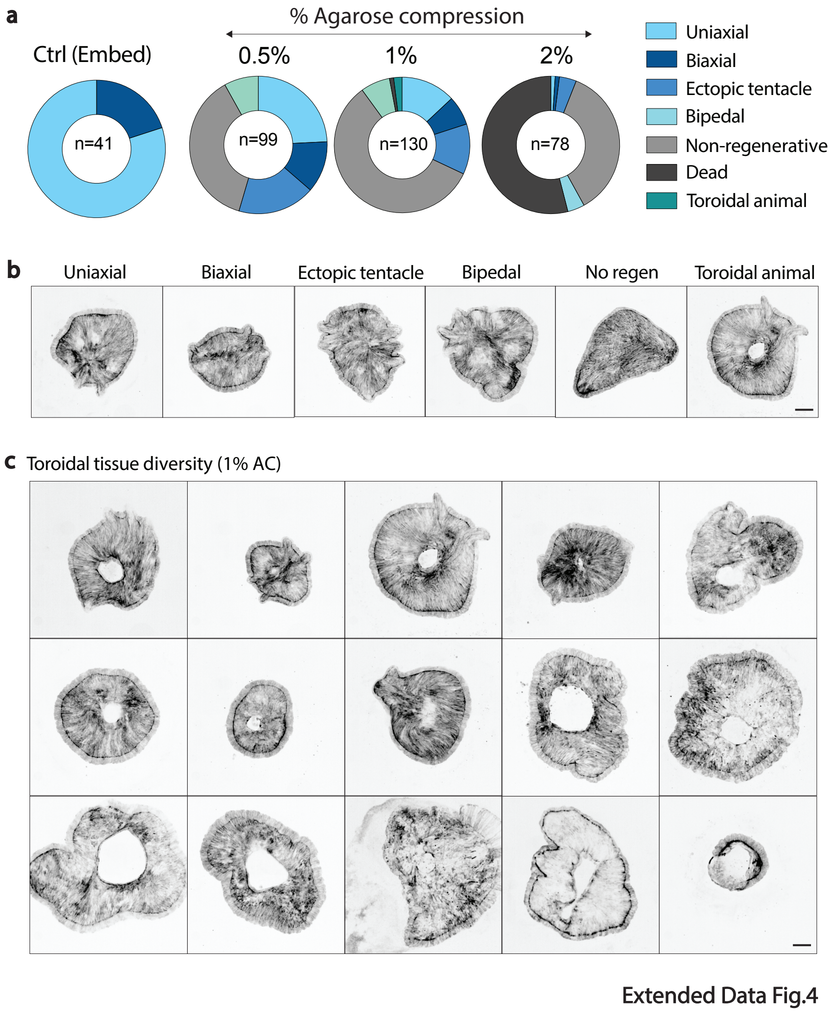


**Extended Data Fig.4 a,** Donut graphs showing phenotype distribution of spheroids at 4dpd under different percentages of AC. **b,** Live spinning disk images **(**Max intensity z-proj) at 4dpd of the various phenotypes observed with regenerating spheroids under compression. Scale bar, 100µm. **c,** Live spinning disk images **(**Max intensity z proj) at 4dpd of toroidal animals obtained with 1% AC of spheroids. **b and c** Scale bars, 100µm.

### I. THEORETICAL DESCRIPTION OF AN ELASTIC ACTIVE NEMATIC SOLID

To study the potential impact of the actin suprastructure on Hydra shape during regeneration, we describe its material properties in terms of an elastic active nematic solid [1, 4, 7]. This is motivated by the mesoglea a gel-like elastic substance [5] that is sandwiched between the animal's two body layers, the (ectodermal) epidermis and (endodermal) gastrodermis. We assume that these are the mechanically most important parts of the animal. We assume that, similar to stress fibres, Hydra's actin cables in the epi- and gastrodermis generate an anisotropic active stress [6]. A possible modulation of Hydra's material properties by genetic or biochemical regulation as well as other parts of the tissue are not part of our description.

We choose a continuum description of the material and the coarse-grained orientation of the active contractile stress is captured by an orientation field $\underline{\hat{p}}$ with $\underline{\hat{p}}^2 = 1$. It is coupled to an order parameter, $S \in [0, 1]$, accounting for the local degree of anisotropy in the active stress. Let us point out that since we assume the orientational order is nematic in character, the sign of $\underline{\hat{p}}$ is inconsequential to the mechanics of the system.

We account for the activity of the elastic material by changes of the material's reference state [3]. In Hydra, such changes result from the internal dynamics of the actin cytoskeleton. Since we confine our analysis to the final states of Hydra regeneration, we impose a time evolution of the reference state, which interpolates between that of a sphere or a torus as the initial stress-free state and a final state incorporating contraction along the orientation field.

To analyse deformations due to changes in the reference state, we introduce an agent-based model. In this model, we partition the volume of the material into polyhedra by means of a three dimensional Voronoi tessellation. The tessellation is such that there are multiple Voronoi cells across the shortest dimensions of the material. In the simulations presented here, this corresponds to the thickness of the material and ensures resistance to bending.

The corresponding Delaunay triangulation describes a network connecting the centres of each Voronoi cell. We account for the average stress through the faces of the Voronoi cells by springs along the edges of the Delaunay network. It should be noted that the Delaunay network is not a representation of any specific structure in the active elastic material. Additionally, the Delaunay network is fixed for the duration of the simulation, thus it does not recreate the dynamics of a change in topology.

We describe changes of the reference state due to growth through time-dependent modifications of the springs' rest lengths. Explicitly, the rest length $l_i$ of spring $i$ at time $t$ with initial orientation $\underline{\hat{l}}_i$ and midpoint position $\underline{r}_i$ is given by

$$l_i = \tilde{l}_i(1 + \zeta(t)) \times \sqrt{1 + \xi(t)S(\underline{r}_i) + \lambda(t)S(\underline{r}_i)[(\underline{\hat{p}} \cdot \underline{\hat{l}}_i)^2 - 0.5]}, \tag{1}$$

where $\tilde{l}_i$ is the initial rest length. The phenomenological parameters $\zeta$, $\xi$ and $\lambda$ represent active strain coefficients. Depending on their sign, they describe relative expansion or contraction. The time-dependence of the strain coefficients reflects the evolution of the reference state. We now discuss their values in turn.

The differential anisotropic strain coefficient $\lambda$ controls changes of the rest length of a connection depending on its alignment with the orientation field, hence induces anisotropic contraction or expansion. We consider linear growth up to a predetermined stall value $\lambda^s$ at time $t^s$:

$$\lambda(t) = \begin{cases} \frac{\lambda^s t}{t^s}, & \text{if } t < t^s. \\ \lambda^s, & \text{otherwise.} \end{cases} \tag{2}$$

We choose $t^s$ and $\lambda$ such that the reference metric evolves slowly and the material remains close to mechanical equilibrium at all times. All results are presented in the long time limit ($t \gg t^s$) where the system is at a steady state.

The differential isotropic strain coefficient $\xi$ controls changes of the rest length depending on the local order parameter, $S$. This then describes isotropic expansion or contraction regardless of the orientation of $\underline{\hat{p}}$ at that point. In the current study we set $\xi = 0$; an in depth discussion of the effect of $\xi$ is available [4]. Finally, the global strain coefficient $\zeta$ represents homogenous, isotropic expansion or contraction, thus only contributes to a global re-scaling of the system. Since we are interested in changes in morphology of the material, we set $\zeta = 0$ for all simulations.

We consider an elastic material, in which the internal dynamics of the nematic field are negligible. Consequently, the values $\underline{\hat{p}} \cdot \underline{\hat{l}}_i$ and $S(\underline{r}_i)$ are fixed at the onset of the simulation. We choose all springs to have the same spring constant, $k = 1$, and evolve the system according to an over-damped Langevin Equation. All simulations run for at least $2t^s$ and the final configurations of the model is shown in figures.

A step in our simulation consists of the following elements: First, each edge in the Delaunay triangulation is given a rest length according to Eq. (1).

The position of each point is then updated according to an over-damped Langevin equation with mobility coefficient

$\mu$, and the total stored elastic energy is given by

$$E = \sum_k \left[ |\underline{r}_k(t) - \underline{r}_j(t)| - l_{kj} \right]^2. \tag{3}$$

Hence the position of point $j$ is updated according to

$$\underline{r}_j(t+1) = \underline{r}_j(t) + \mu \sum_k \left[ |\underline{r}_k(t) - \underline{r}_j(t)| - l_{kj} \right] \hat{\underline{r}}_{kj}(t), \tag{4}$$

where the points $\underline{r}_k$ are those linked to point $\underline{r}_j$ by an edge of the Delaunay triangulation, with rest length $l_{kj}$ and $\hat{\underline{r}}_{kj}$ is a unit vector pointing from point $k$ to point $j$. In all data presented $\mu = 0.05$. We simulate for $T = 10^5$ time steps with $t^s = T/2$.

This completes the description of our agent-based model.

##### A. Actin suprastructure in Hydra ectoderm and endoderm

In our simulations, we consider a single closed surface with finite thickness, roughly mimicking the structure of Hydra. Furthermore, we take the orientation field $\hat{\underline{p}}$ to be tangential to the surface, similar to the orientation of actin cables in Hydra. We recall that the resolution of the simulation is chosen to ensure multiple Voronoi cells over the thickness of the surface to ensure resistance to bending.

Hydra consists of two body layers, the outer ectoderm and inner endoderm. Super-cellular/Supra-cellular actin cables are present in both layers of the animal and are perpendicular to each other. That is to say that if $\hat{\underline{p}}$ describes the orientation of the cables in the ectoderm, then we can define a second vector field $\hat{\underline{q}}$, such that $\hat{\underline{p}}.\hat{\underline{q}} = 0$, which describes the orientation of the actin cables in the endoderm.

Since $\hat{\underline{p}}$ and $\hat{\underline{q}}$ are unit vectors we can write

$$\hat{\underline{p}} = \cos(\psi)\hat{\underline{x}} + \sin(\psi)\hat{\underline{y}} \tag{5}$$

$$\hat{\underline{q}} = \cos(\psi \pm \pi/2)\hat{\underline{x}} + \sin(\psi \pm \pi/2)\hat{\underline{y}}, \tag{6}$$

where $\psi$ indicates the in-plane orientation of $\hat{\underline{p}}$ and $\hat{\underline{x}}$ and $\hat{\underline{y}}$ denote two orthogonal unit vectors in the surface's tangent plane.

When transforming between $\hat{\underline{p}}$ and $\hat{\underline{q}}$ the position and charge of topological defects is preserved, however the phase changes by $\pi/2$. For defects with charge different from 1, this indicates a rotation of the defect. For $+1$ topological defects a phase change of $\pi/2$ corresponds to a change between asters and vortices, and in the case of spirals to a change of the chirality.

The behaviour of the two layered system is completely captured by the introduced model if the order parameters of both layers is the same. Indeed, since the field $\hat{\underline{q}}$ is completely specified by the field $\hat{\underline{p}}$, it does not introduce any additional degrees of freedom. Explicitly, consider Eq. 1. The orientation field enters in the following term

$$\lambda(t)S(\underline{r}_i)[(\hat{\underline{p}} \cdot \hat{\underline{l}}_i)^2 - 0.5]. \tag{7}$$

Since $\hat{\underline{p}}$ and $\hat{\underline{l}}$ are unit vectors we can write this as

$$\lambda(t)S(\underline{r}_i)[\cos^2(\beta) - 0.5]. \tag{8}$$

where $\beta$ is the angle between $\hat{\underline{p}}$ and $\hat{\underline{l}}$. For the second vector field $\hat{\underline{q}}$, we have analogously

$$\lambda_2(t)S(\underline{r}_i)[(\hat{\underline{q}} \cdot \hat{\underline{l}}_i)^2 - 0.5]. \tag{9}$$

Once again we can write

$$(\hat{\underline{q}} \cdot \hat{\underline{l}}_i)^2 - 0.5 = \cos^2(\beta \pm \pi/2) - 0.5 \tag{10}$$

$$= \sin^2(\beta) - 0.5 \tag{11}$$

$$= 0.5 - \cos^2(\beta) \tag{12}$$

$$= -[(\hat{\underline{p}} \cdot \hat{\underline{l}}_i)^2 - 0.5]. \tag{13}$$

Here, we have employed the angle addition formula for cos and the Pythagorean identity. Thus the sum of Eqs. 7&9 can be written as

$$[\lambda(t) - \lambda_2(t)]\, S(\underline{r}_i)[(\hat{\underline{p}} \cdot \hat{\underline{l}}_i)^2 - 0.5]. \tag{14}$$

In summary, by introducing the field $\hat{\underline{q}}$ into Eq. 1, we have merely changed the interpretation of the phenomenological parameter $\lambda$, which now is interpreted as the combination of extensions and contractions in both the endoderm and ectoderm. For this reason it is sufficient to consider in our simulations a single vector field $\hat{\underline{p}}$.

#### II. SIMULATION OF SPHERICAL SHELLS

All simulations performed in this paper are for thin active elastic materials, in which the thickness is much smaller than the other dimensions. We refer to these as shells. For all visualisations, the mid-plane of the thin material is approximated by coarse-graining the point cloud to a set of evenly distributed bins in the plane shell. This mid-plane of the shell is what is displayed in the images of the final configuration in all figures.

##### A. Preparation of the Voronoi tessellation

When simulating an initially spherical shell, the simulation is prepared as follows. First, the shell thickness is designated $t$ in units of the spherical inner radius. Thus the volume of the simulated material is given by $V = 4\pi((1+t)^3 - 1)/3$. In this volume $N$ points are randomly placed with uniform probability density. Finally, a single additional auxiliary point is placed at the origin, shown in red in Fig. 1a. We then generate a Delaunay triangulation from these points. After removing the auxiliary point at the origin, there are no connections across the body of the sphere.

Simulations originating from spherical shells contain 2000 points. Spherical shells are initialized with an inner radius of $R = 1$ and thickness $R/10$. We set $\lambda = 1$.

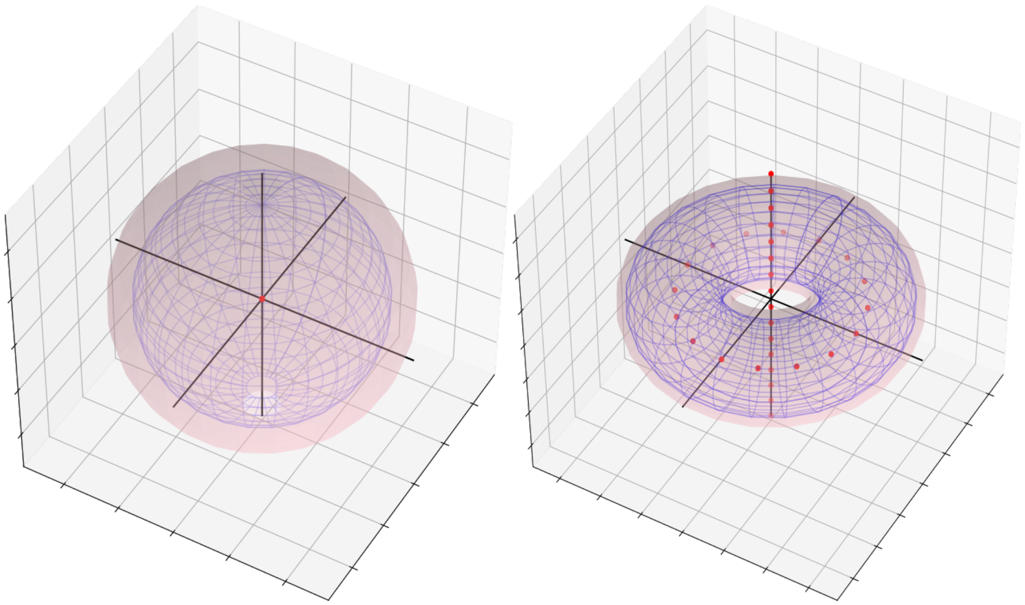

FIG. 1. Schematic of the initialisation of (a) spherical and (b) toroidal shells. The blue wireframe shows the inner surface and the pink sphere shows the outer surface. Auxiliary points are shown in red.

##### B. Compression of the active material

External compression of the material is captured by an additional elastic force on each node of the Delaunay triangulation that are sufficiently far from the origin of the coordinate system in the compression direction, $\hat{\underline{c}}$. The origin of the coordinate system is centred on the centre of mass of the material. The additional restorative force experienced by particle $j$ is given by

$$\underline{f}_j = \begin{cases} -k_c(\underline{r}_j.\hat{\underline{c}} - R_c)\hat{\underline{c}}, & \underline{r}_j.\hat{\underline{c}} > R_c \\ 0, & |\underline{r}_j.\hat{\underline{c}}| < R_c \\ k_c(\underline{r}_j.\hat{\underline{c}} - R_c)\hat{\underline{c}} & \underline{r}_j.\hat{\underline{c}}, < -R_c \end{cases} . \tag{15}$$

In our simulations, we set $R_c$ to be $1.5R$, where $R$ is the initial inner radius of the spherical shell, and $k_c = 1$. Compression only has a significant effect on the outcome of the simulation in the case of a non-regenerating Hydra on which it causes the +1 topological defects to invert. Compression is not included in the toroidal simulations since they correspond to Hydra post release.

##### C. Designation of the director field

We assume that the director field is constant across the thickness of the surface. This is described at the start of a simulation as a function of the angular position on the surface of the sphere given by the polar and azimuthal angles $\Theta$ and $\Phi$, respectively. The director field at each point on the sphere can then be expressed by the angle it makes with the meridians of the sphere, $\psi$. This gives

$$\hat{\underline{p}} = \cos(\psi)\hat{\underline{g}}_\theta + \sin(\psi)\hat{\underline{g}}_\phi, \tag{16}$$

where $\hat{\underline{g}}$ are the orthonormal basis vectors on the surface of the sphere.

We calculate the orientation of the nematic field at each point using a stereographic projection to place these defects onto the complex plane $z(\Theta, \Phi) = R\cot(\Theta/2)e^{i\Phi}$, where $R$ is the radius of the sphere. We complete the nematic field by minimising the Frank free energy.

The in-plane nematic director that minimises the Frank free energy around a set of $j$ defects with positions $z_j$ and charges $m_j$ is given by $\hat{\underline{n}} = (\cos(\alpha), \sin(\alpha))$ with

$$\alpha = \alpha_0 + \sum_j \mathrm{Im}(\ln(z - z_j)^{m_j}). \tag{17}$$

Here we have introduced a global phase $\psi_0$. The director field is then projected back onto the sphere using $\psi(\Theta, \Phi) = \Phi - \alpha(\Theta, \Phi)$. This provides a nematic field that minimizes the elastic energy around the defects specified in the one constant approximation. Care must be taken to ensure $\sum_j m_j = 2$ consistent with the Poincaré Hopf Theorem [2].

The order parameter associated with the director field is given by

$$S = 1 - \sum_j \exp(-\Delta_j/\epsilon) \tag{18}$$

where $\Delta_j$ is the geodesic distance from defect $j$ and we have introduced the defect core radius $\epsilon$.

A summary of the topological defects in each simulation is included in tables. I-V.

For completeness, we also include a spherical surface with topological defects associated with the tentacles on the Hydra. In keeping with the topological constraints, each tentacle has a +1 defect at its tip and a pair of −1/2 defects at its base. This makes them each topologically neutral and thus the total number of tentacles is not topologically constrained.

TABLE I. **Sphere with two +1 defects, compression perpendicular to the axis passing through the poles: Fig. 2C (Main text)**

| Defect charge | Feature | $\Theta$ | $\Phi$ |
|---|---|---|---|
| +1 | Mouth | 0 | 0 |
| +1 | Foot | $\pi$ | 0 |
| - | Compression direction | $\pi/2$ | $\pi/2$ |

TABLE II. **Sphere with three +1 defects, compression perpendicular to the plane containing the +1 defects: Fig. 2C (Main text)**

| Defect charge | Feature | $\Theta$ | $\Phi$ |
|---|---|---|---|
| +1 | Mouth 1 | $\pi/3$ | 0 |
| +1 | Mouth 2 | $\pi/3$ | $\pi$ |
| +1 | Foot | $\pi$ | 0 |
| -1/2 | Additional defect 1 | $\pi/6$ | $3\pi/2$ |
| -1/2 | Additional defect 2 | $\pi/6$ | $\pi/2$ |
| - | Compression direction | $\pi/2$ | $\pi/2$ |

TABLE III. **Sphere with two +1 defects, compression along the axis passing through the poles: Fig. 4a (Main text)**

| Defect charge | Feature | $\Theta$ | $\Phi$ |
|---|---|---|---|
| +1 | Mouth 1 | 0 | 0 |
| +1 | Foot | $\pi$ | 0 |
| - | Compression direction | 0 | 0 |

#### III. SIMULATION OF TOROIDAL SHELLS

##### A. Preparation of the Voronoi tessellation

When simulating an active elastic material with a toroidal topology, the simulation is prepared as follows. First, the shell thickness is designated $t$ in units of the inner, minor radius of the torus. Thus the volume of the simulated material is given by $V = 2\pi^2((1+t)^2 - 1)$. In this volume $N$ points are randomly placed with uniform probability density.

Due to the more complex topology of the torus, a set of additional auxiliary points are placed in and around the torus. These are located on a ring inside the torus, and on a line that threads the centre of the torus, Fig. 1b. We then generate a Delaunay triangulation including these points. The auxiliary points in and around the torus are removed before the simulation begins and thus ensure that the hollow regions of the simulation are not connected. Toroidal simulations are initialised with 10000 points, an inner minor radius of $R = 1$, and a thickness of $R/10$. We set $\lambda = 1.5$.

##### B. Designation of the director field on the torus

We assume that the director field is constant across the thickness of the material, such that it can be described as a function of the position on the surface of the torus given by the curvilinear coordinates $\Theta$ and $\Phi$, which designate the

TABLE IV. **Sphere with four +1 defects, compression perpendicular to the plane containing the +1 defects: SI Fig.2**

| Defect charge | Feature | $\Theta$ | $\Phi$ |
|---|---|---|---|
| +1 | Mouth | 0 | 0 |
| +1 | Foot | $\pi$ | 0 |
| +1 | Tentacle 1 tip | $2\pi/8$ | 0 |
| -1/2 | Tentacle 1 base | $3\pi/8$ | $\pi/3$ |
| -1/2 | Tentacle 1 base | $3\pi/8$ | $5\pi/3$ |
| +1 | Tentacle 2 tip | $2\pi/8$ | $\pi$ |
| -1/2 | Tentacle 2 base | $3\pi/8$ | $2\pi/3$ |
| -1/2 | Tentacle 2 base | $3\pi/8$ | $4\pi/3$ |
| - | Compression direction | $\pi/2$ | $\pi/2$ |

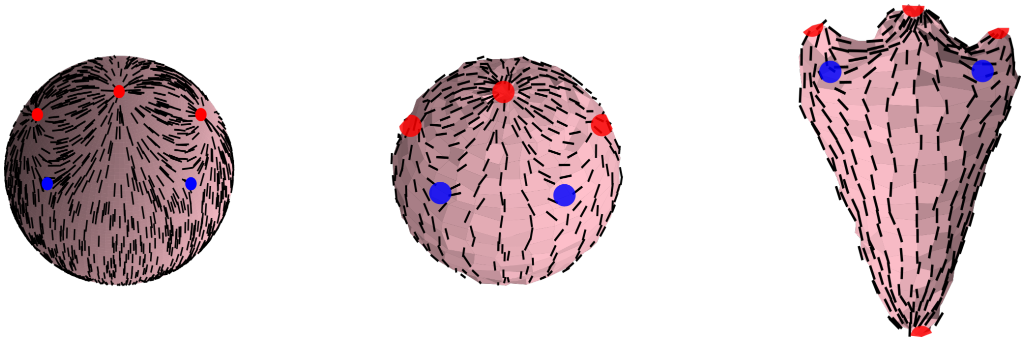

FIG. 2. Surface evolving with the topological defects associated with a Hydra featuring two tentacles. Left) Nematic texture around the specified defects. Middle) Starting point of the simulation. Right) End point of the simulation.

TABLE V. **Torus with two +1 defects: Fig.4c (Main text)**

| Defect charge | Feature | $\Theta$ | $\Phi$ |
|---|---|---|---|
| +1 | Mouth | 0 | 0 |
| +1 | Foot | 0 | $\pi$ |
| -1/2 | Additional defect 1 | 0 | $3\pi/2$ |
| -1/2 | Additional defect 2 | 0 | $\pi/2$ |
| -1/2 | Additional defect 3 | $\pi$ | 0 |
| -1/2 | Additional defect 4 | $\pi$ | $\pi$ |

poloidal and toroidal directions, respectively. Without loss of generality we assign $\Theta = 0$ to denote the outer equator of the torus.

The director field at each point on the sphere can then be given by the angle it makes with the meridians of the torus, $\psi$. This gives

$$\underline{\hat{p}} = \cos(\psi)\underline{\hat{g}}_\theta + \sin(\psi)\underline{\hat{g}}_\phi \tag{19}$$

Where $\underline{\hat{g}}$ are the orthonormal basis vectors on the surface of the torus.

For simulations corresponding to non-regenerating toroidal Hydra, we set $\psi = 0$ everywhere as is observed in experiments, such that it features no defects.

For simulations corresponding to toroidal Hydra with delayed regeneration, we use a director field with defects. To generate the orientation field around a set of defects, first we generate the field on flat surface with periodic boundaries. The director field is given by

$$\psi = \sum_j k_j \phi_j, \tag{20}$$

where $k_j$ is the charge and $\phi_j$ is the polar angle relative to defect $j$, respectively. A smoothly varying phase may be added to this field to ensure the desired phase of the defect cores.

The order parameter associated with the director field is calculated as

$$S = 1 - \sum_j \exp(-\Delta_j/\epsilon), \tag{21}$$

where $\Delta_j$ is the Euclidian distance from defect $j$ and we have introduced the defect core radius $\epsilon$.

Finally, the Landau-de Gennes free energy of the field is minimised iteratively in regions for which $S > S_c$. This cut-off for $S$ is to fix the position and orientation of the defects, otherwise they would migrate during energy minimisation.

The defect positions for recovering toroidal hydra are given in Table. V. The non-recovering toroidal hydra features no topological defects.

[1] Daniel Duffy and John S Biggins. Defective nematogenesis: Gauss curvature in programmable shape-responsive sheets with topological defects. Soft Matter, 16(48):10935–10945, 2020.

[2] Diana Khoromskaia and Gareth P Alexander. Vortex formation and dynamics of defects in active nematic shells. New Journal of Physics, 19(10):103043, 2017.

[3] DA Matoz-Fernandez, Fordyce A Davidson, Nicola R Stanley-Wall, and Rastko Sknepnek. Wrinkle patterns in active viscoelastic thin sheets. Physical Review Research, 2(1):013165, 2020.

[4] DJG Pearce, S Gat, G Livne, A Bernheim-Groswasser, and K Kruse. Defect-driven shape transitions in elastic active nematic shells. arXiv preprint arXiv:2010.13141, 2020.

[5] Hiroshi Shimizu, Roland Aufschnaiter, Li Li, Michael P Sarras Jr, Dorin-Bogdan Borza, Dale R Abrahamson, Yoshikazu Sado, and Xiaoming Zhang. The extracellular matrix of hydra is a porous sheet and contains type iv collagen. Zoology, 111(5):410–418, 2008.

[6] Anniek Stokkermans, Aditi Chakrabarti, Kaushikaram Subramanian, Ling Wang, Sifan Yin, Prachiti Moghe, Petrus Steenbergen, Gregor Mönke, Takashi Hiiragi, Robert Prevedel, et al. Muscular hydraulics drive larva-polyp morphogenesis. Current Biology, 32(21):4707–4718, 2022.

[7] M. Warner and E.M. Terentjev. Liquid Crystal Elastomers. International Series of Monographs on Physics. OUP Oxford, 2007.
